# Parasitism effects on coexistence and stability within simple trophic modules

**DOI:** 10.1101/289819

**Authors:** Loïc Prosnier, Vincent Médoc, Nicolas Loeuille

**Affiliations:** Sorbonne Université, Université Paris Diderot, Université Paris-Est Créteil, CNRS, INRA, IRD, institute of Ecology and Environmental Science - Paris (iEES-Paris), Campus Pierre et Marie Curie, 4 place Jussieu, 75005 Paris, France; Equipe Neuro-Ethologie Sensorielle, ENES/Neuro-PSI, CNRS UMR 9197, Université de Lyon/Saint-Etienne, 23 rue Michelon, 42023 Saint-Etienne Cedex 2, France

**Keywords:** apparent competition, direct competition, host-parasite interaction, interaction effect, paradox of enrichment, prey-predator interaction, stability, virulence

## Abstract

Parasites are important components of food webs. Although their direct effects on hosts are well-studied, indirect impacts on trophic networks, thus on non-host species, remain unclear.

In this study, we investigate the consequences of parasitism on coexistence and stability within a simple trophic module: one predator consuming two prey species in competition. We test how such effects depend on the infected species (prey or predator). We account for two effects of parasitism: the virulence effect (parasites affect the infected species intrinsic growth rate through direct effects on fecundity or mortality) and the interaction effect (increased vulnerability of infected prey or increased food intake of infected predators).

Results show that coexistence is favored when effects have intermediate intensity. We link this result to modifications of direct and apparent competitions among prey species. Given a prey infection, accounting for susceptible-infected population structure highlights that coexistence may also be reduced due to predator-parasite competition.

Parasites affect stability by modulating energy transfer from prey to predator. Predator infection therefore has a stabilizing effect due to increased energy fluxes and/or predator mortality.

Our results suggest that parasites potentially increase species coexistence. Precise predictions however require an assessment of various parasite effects. We discuss the implications of our results for the functioning of trophic networks and the evolution of foraging strategies within food webs.

## 1. Introduction

Many studies on food webs show that parasites are omnipresent, with a high biomass (Kuris et al., 2008) and making a large proportion of antagonistic interactions (Amundsen et al., 2009; Hudson, Dobson, & Lafferty, 2006). Although parasites are expected to have large impacts on diversity and stability (Poulin, 2010; Wood & Johnson, 2015), exact consequences appear difficult to estimate due to complexity of ecological networks and the diversity of parasite effects (Hatcher, Dick, & Dunn, 2006, 2014; Welch & Harwood, 2011). We therefore need an integrative perspective on the effects of parasitism in multi-species systems (ecosystem parasitology, Hatcher & Dunn, 2011; Tompkins, Dunn, Smith, & Telfer, 2011). In the present work, we analyze the consequences of parasitism on coexistence and stability, using a trophic module approach.

We investigate two effects of parasites, called hereafter “virulence effects” and “interaction effects”. Virulence effects embody the direct consequences of infection parasites typically reduce the fecundity and/or survival of their hosts (Coors & De Meester, 2011), thereby impacting their intrinsic growth rates. Such virulence effects are well-documented: Decaestecker, Vergote, Ebert, & De Meester (2003) for instance showed reduced fecundity and increased mortality of *Daphnia magna* when infected by bacteria (*Pasteuria*) or fungi (Microsporidia). Similar effects of trematodes on *Daphnia obtusa* have been observed by Schwartz & Cameron (1993). Such virulence effects may propagate at the population level, decreasing host biomass and affecting competitive hierarchies among species (Decaestecker, Verreydt, De Meester, & Declerck, 2015).

By affecting the phenotype of their hosts, parasites may also change the trophic interactions involving these hosts in the network (hereafter, “interaction effects”). This is well established for trophically-transmitted parasites as natural selection on the parasite may affect its host appearance or behavior (such modifications can then be seen as an extended phenotype of the parasite) in a way that increases its vulnerability to predation (trophic manipulation), thereby facilitating transmission (Butler IV, Tiggelaar II, Shields, & Butler V, 2014; Cézilly, Thomas, Médoc, & Perrot-Minnot, 2010; Jacquin, Mori, & Médoc, 2013; Lefèvre et al., 2009). However, modifications of predator-prey interactions may also be simply an indirect effect of parasitism happening in the absence of trophic transmission (Duffy, Hall, Tessier, & Huebner, 2005; Hudson, Dobson, & Newborn, 1992; Peterson & Page, 1988). For instance, daphnia infected by *Pasteuria ramosa* have a red coloration that makes them more catchable by *Anisop* (Goren & Ben-Ami, 2017). An infected predator may also increase its food intake (predation rate) to compensate the energetic costs incurred by the infection (Bernot & Lamberti, 2008; Dick et al., 2010; Lettini & Sukhdeo, 2010).

We here assess such consequences, defining coexistence as the possibility of maintaining all species (predators and prey species) and stability based on the occurrence of oscillating population dynamics. In our system, coexistence among the two prey species depends on the balance between direct (i.e. resource based) and apparent competition (i.e. competition mediated by the predator presence, Holt [1977]). As illustrated by classical experiments (Gause, 1934), direct competition is an important constraint for coexistence. The inclusion of a predator in a competitive system would affect coexistence through apparent competition (Holt, 1977). In such situations, one prey negatively affects the other prey by increasing the predator population. By combining the two competitions (direct and apparent competition), coexistence is allowed when the most competitive species is also the most vulnerable prey (Holt, Grover, & Tilman, 1994). Consequently, coexistence requires a balance between direct and apparent competition that may be affected by parasites. Virulence effects may for instance reduce competitive ability of the host species. Competition between *Daphnia magna* and *D. pulex*, usually favoring *D. magna*, may be reversed when *D. magna* are infected by microsporidian and bacteria (Decaestecker et al., 2015). Now consider parasitism on the predator. By decreasing predator density, a parasite with virulent effects may decrease apparent competition and thereby favor the preferred prey species, which is also the best direct competitor. Therefore, in prey infection as in predator infection, parasites with virulent effects likely affect coexistence by changing the relative intensity of the two types of competition (Fig. 1b). Interaction effects may be equally important. When they modify predation, parasites directly alter apparent competition. An increased predation on the best direct competitor, should favor the other prey species. Such parasite-induced modification of predation by snails (*Littorina littorea)* on ephemeral macroalgae, for instance, modify biomass and composition of intertidal communities (Wood et al., 2007). Under such scenarios, parasites favor coexistence at intermediate effects, while extremely high or low effects decrease coexistence by altering the balance between the two types of competition (Fig. 1b).

**Figure 1.**
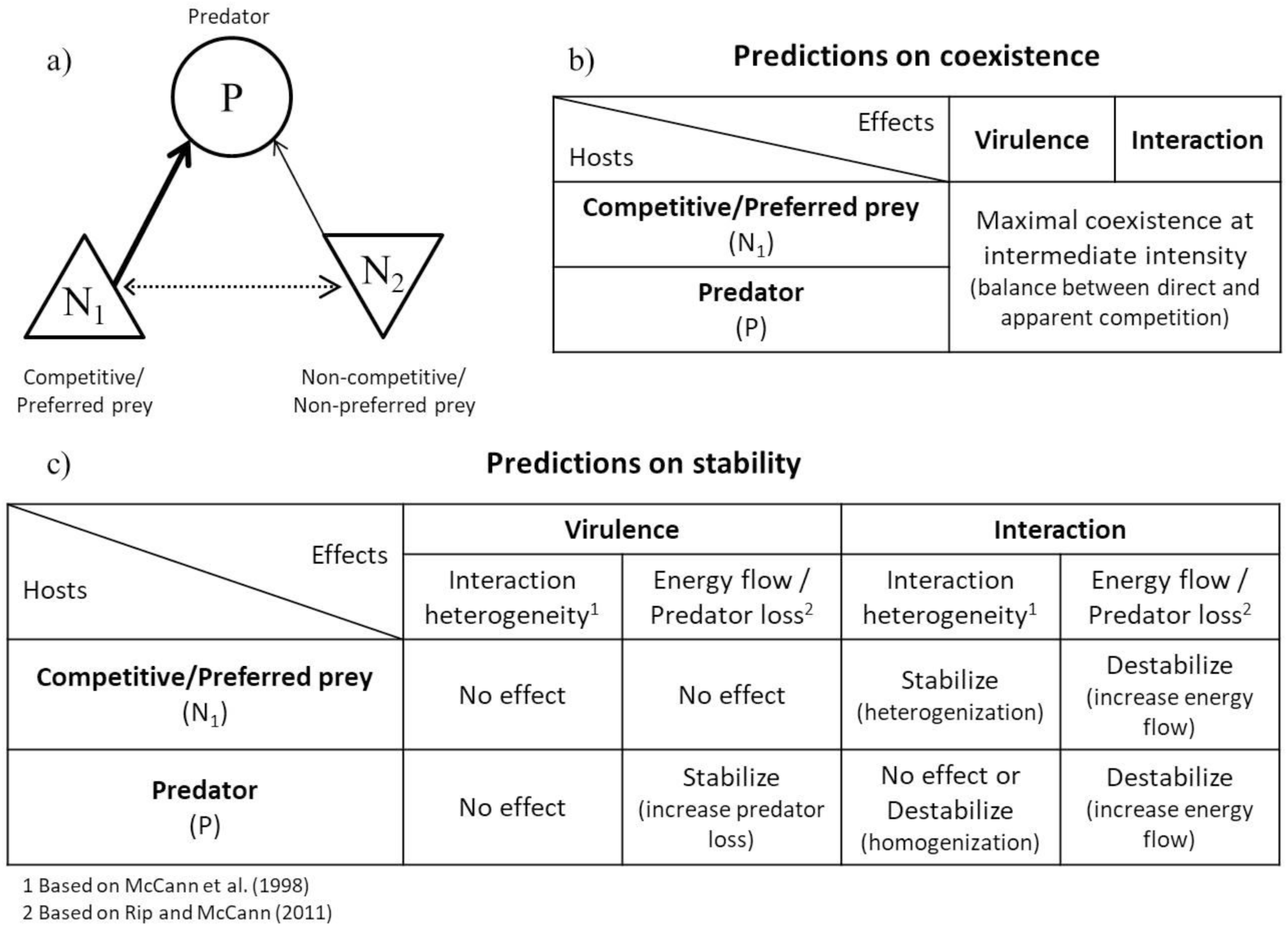
Presentation of the trophic module and predictions. a) The module before infection consists of *P*, the predator, *N_1_*, the competitive/preferred prey, and N_2_, the non-competitive/non-preferred prey; solid arrow, the predation, dashed arrow, the direct competition. Predictions on how coexistence (Table b) and stability (Table c) depend on the infection scenario (identity of species infected, virulence or interaction effects).

We also analyze the effects of parasites on the stability of the system (Fig. 1c). Considering the paradox of enrichment, a classical result in ecological theory is that stability decrease with increase of the relative intensity of energy flow (predation rate) versus predator loss rate (predator mortality rate) (Rip & McCann, 2011; Rosenzweig, 1971). Consider virulence effects. For an infected prey, we do not expect effects on stability as neither predation rate nor predator mortality rate is modified. For an infected predator, however, infection increases the predator mortality rate and thus stabilizes the system. Now consider the interaction effect. As it directly increases the predation rate, it is expected to destabilize the system.

Next to such “energy flow” aspects, classical ecological theory also suggests that stability is enhanced when weak and strong interactions coexist within a trophic module (McCann, Hastings, & Huxel, 1998). Heterogeneous systems made of few strong links and many weak links are more stable than homogeneous systems. Such stability constraints are not affected by virulence effects as they do not change the balance of interactions. On the contrary, interaction effects directly modify the distribution of interaction strengths and increasing predation on the most consumed prey should increase heterogeneity, and thus stability. When the predator is infected, the changes in the distribution of interaction strengths depend on how the parasite affects interaction rates. If all interaction rates increase in the same proportion stability should not be modified. When all rates increase by a given, fixed change, interaction strengths should be homogenized and the system destabilized.

Many studies of parasite effects focus on one species (for instance describing virulence effects) or on a few species in interactions (for instance describing trophic interaction modifications by parasites). Nevertheless, the review by Hatcher et al. (2006) shows how the consequences of parasitism may extend to more complex systems. Here, we tackle the effects of parasitism on coexistence and stability, explicitly considering a predation-competition context (Fig. 1a). Using such a system allows us to consider “parasite-modified competition [and interactions] with apparent competition” as suggested by Hatcher et al. (2006).

We first consider infection of the prey, then of the predator species. In each case, we first tackle virulence effects (on reproduction or mortality rates), then interaction effects (changes in trophic interactions). To allow a more tractable analysis, we first simplify the system, by considering that parasite effects are simple modifications of the host parameters. Such an approach is however limited, as it neglects important ecological feedbacks (e.g. parasite-predator competition when prey species are infected). Therefore, as a second step, we consider a system in which the host population is structured in susceptible and infected individuals. Our aims are to understand how the consequences of parasitism depend on the host trophic level or on parasite effect (virulence or interaction). We expect that prey parasitism increases coexistence (i.e. presence of the three species) when the best competitor is infected, as the parasite then decreases direct competition while increasing apparent competition. When the predator is infected, an intermediate level of parasitism is expected to favor coexistence (Fig. 1b). Concerning stability, we predict that virulence effects will not change stability, except when infected predators undergo large mortality rates (the system should then be stabilized) (Fig. 1c). Predicting how interaction effects alter stability is more difficult as they may modify both the energy transfer (destabilizing the system), and the distribution of interaction strengths within the module (Fig. 1c).

## 2. Model & Methods

### 2.1. General approach

To study the effects of parasites on a predation-competition system we proceed in two steps. First, we use an unstructured model in which the parasite dynamics are not explicitly included. We instead assume that parasite effects can be modeled by simple variations in the parameters of the host population dynamics.

We then explicit parasite dynamics by structuring the host population in susceptible and infected individuals, as in Anderson & May (1986). Under such scenarios, an explicit competition between the parasite and the predator takes place under prey infection scenarios. It therefore gives a more complete account of the feedbacks that occur between the parasite and the trophic module.

### 2.2. Presentation of the unstructured system

We rely on the two prey-one predator model analyzed by Hutson & Vickers (1983), so that local and global stability conditions are already known. The model considers both intra and interspecific competition for the prey species and a linear functional response for the trophic interaction:

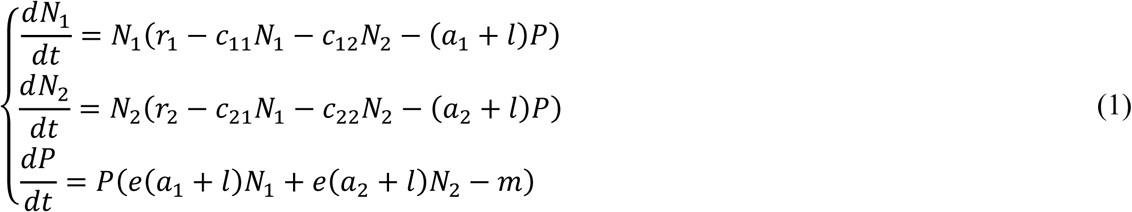

with *N*_*i*_ the density of the prey species *i*, *P* the predator density, *r*_*i*_ the intrinsic growth rate of prey species *i*, *c*_*ii*_ its *per capita* intraspecific competition rate, *c*_*ij*_ the *per capita* effect of interspecific competition of species *j* on species *i*, *a*_*i*_ the attack rate on species *i*, *e* the conversion efficiency, *m* the predator intrinsic mortality rate and *l* the increased food requirement of infected predators. Parameter biological interpretation, dimensions and default values are given in Table 1.

**Table 1.**
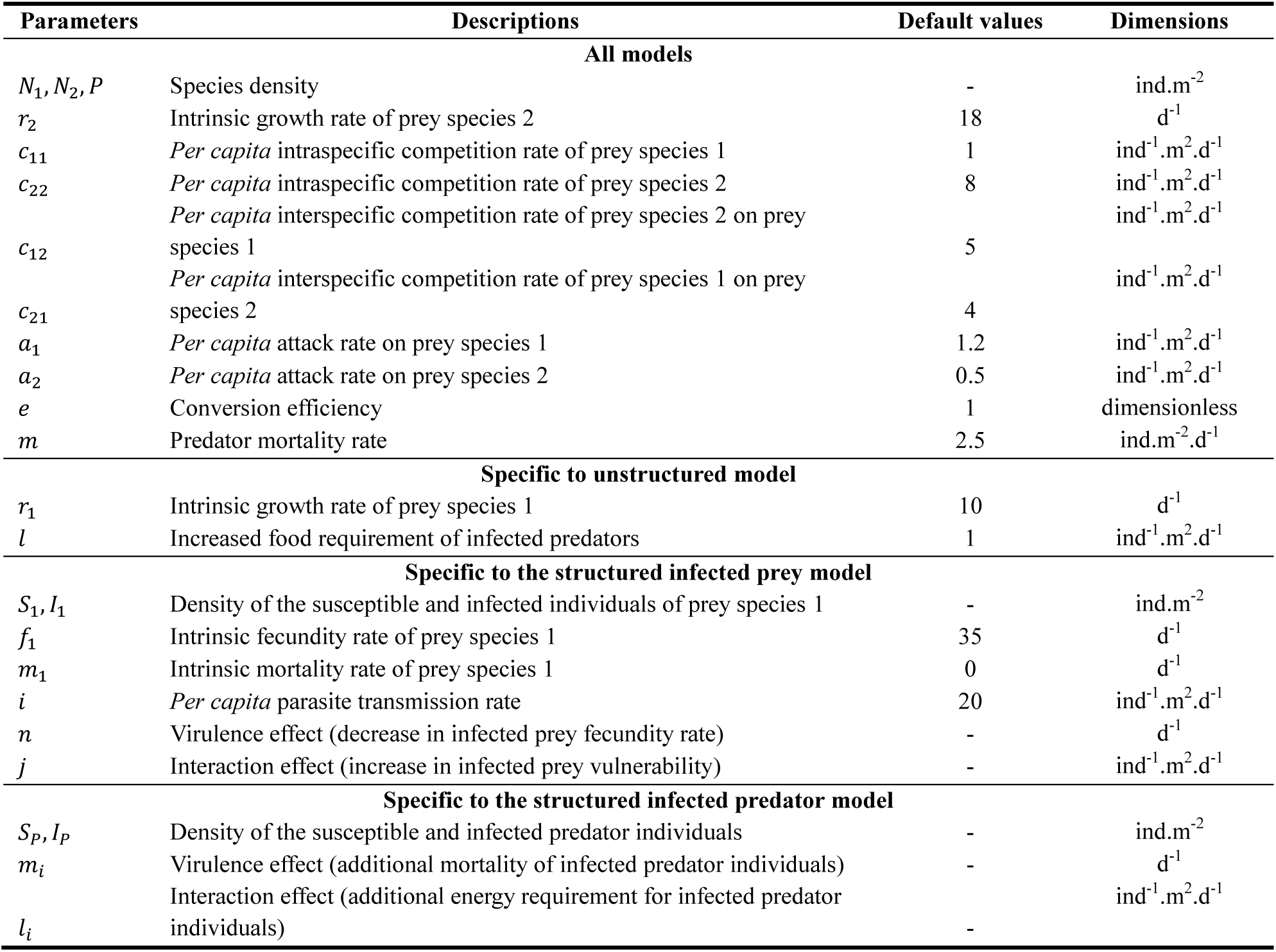
Model parameters (as well as their default values) and variables (default values are based on values proposed in Hutson & Vickers [1983]).

Using system (1), we mimic the two effects (virulence effect and interaction effect) of the parasite. In case of prey infection (on *N*_1_), the virulence effect is modeled through a decreased growth rate (*r*_1_) and the interaction effect through an increased predation on infected hosts (*a*_1_). In case of predator infection, the virulence effect is modeled through an increase of mortality rate (*m*) and the interaction effect by a simultaneous increase of the two attack rates (*a*_1_ and *a*_2_).

### 2.3. Presentation of the Susceptible-Infected structured systems

#### 2.3.1. Structured model of prey infection

We now include infected prey population structure in the initial model (Eq. (1)) through a SI-structured model (Anderson & May, 1986; Kermack & McKendrick, 1927):

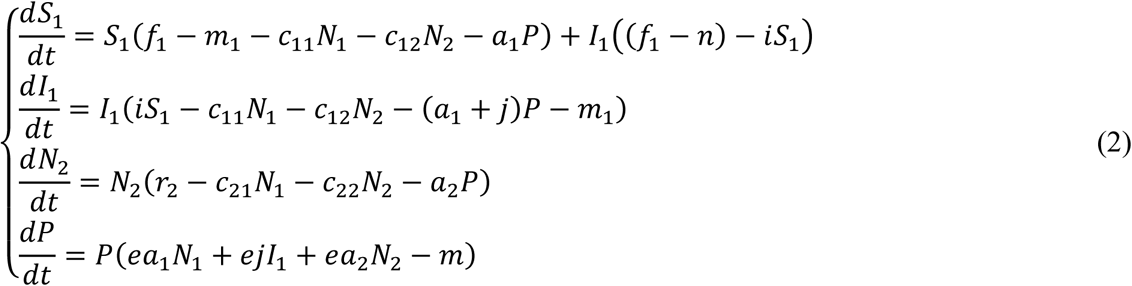

with *S*_1_ and *I*_1_ the susceptible and infected prey densities (*N*_1_ = *S*_1_ + *I*_1_), *f*_1_ its intrinsic fecundity rate, *m*_1_ its intrinsic mortality rate and *i* the *per capita* parasite transmission rate.

In this model, virulence effects are modeled through a reduction of fecundity (parameter *n*) while interaction effects are modeled through changes in prey vulnerability (parameter *j*).

#### 2.3.2. Structured model of predator infection

We similarly consider a structured model in which predators are infected. The initial model (Eq. (1)) can then be rewritten:

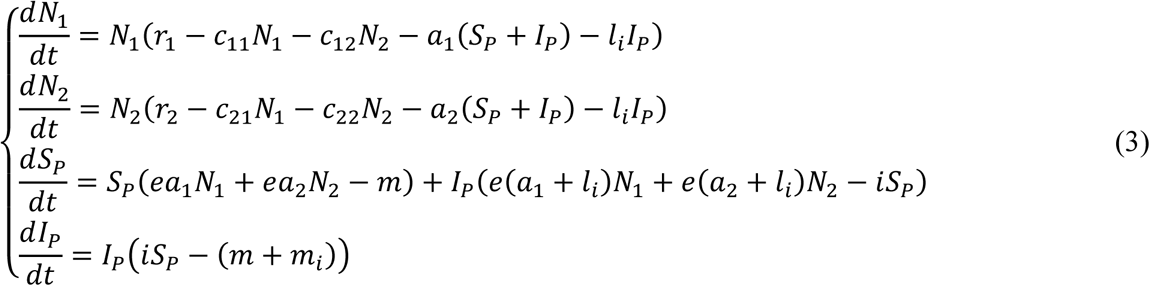

with *S*_*P*_ and *I*_*P*_ the densities of susceptible and infected predators (*P* = *S*_*P*_ + *I*_*P*_).

Virulence effects are considered through an increase in mortality rate (parameter *m*_*i*_) while interaction effects modify the predation rate (parameter *l*_*i*_), assuming that infected predators have larger energetic requirements.

### 2.4. Method of analysis of the different models

We systematically explored the consequences of the two effects of the parasite. We analyzed how they alter the coexistence of the three species (in the unstructured model) and of the four species (including the parasite) in the structured models. We then analyzed their consequences for stability by investigating the type of dynamics (stable point, cycles) occurring under different parasitism scenarios.

Our analysis relies on a number of assumptions. First, we assume that before infection, the predator has a greater attack rate on the most competitive prey (prey species 1). Therefore, the presence of the predator facilitates coexistence among prey species through apparent competition (Holt et al., 1994). Such assumptions lead to the following parameter constraints: *a*_1_ > *a*_2_, *r*_1_*c*_22_ > *r*_2_*c*_12_ and *r*_1_*c*_21_ > *r*_2_*c*_11_ (Hutson & Vickers, 1983). To emphasize the role of predation, we focus on cases where interspecific competition is dominant (*c*_11_*c*_22_ < *c*_12_*c*_21_) so that the two competitors cannot coexist in the absence of predators (Hutson & Vickers, 1983). Note that such constraints only apply before infection. The different scenarios of parasitism and the intensity of parasite effects indeed affect prey competitive abilities (when prey species are infected) and trophic interaction rates (through interaction effects). In scenarios of prey infection, we consider that the host species is the most competitive species (i.e. species 1).

For all scenarios, we perform numerical analyses using Mathematica® 11.1.1 (Wolfram research). First, using the structured models, we simulate the effect of a parasite addition in a non-infected system. Such simulations illustrate how the impacts vary depending on the parasitism effect (virulence or interaction) and on the infected species (predator or prey). For both the unstructured and the structured models, we then analyze the effects of parasitism more globally, through 2D-bifurcation diagrams (one dimension showing variations in virulence effects, the other variations in interaction effects).

## 3. Results

### 3.1. Effects of parasite addition in the three-species structured models

Considering the high variety of expected effects of parasitism (Figs 1b-c), we start by presenting some examples of possible effects of adding a parasite on the coexistence and the stability of the three-species system. First note that virulence effects may increase coexistence because they reduce the competitive ability of the most competitive species. For instance, in Fig. 2a, parasitism on the most competitive species (species 1) eventually allows the invasion of the predator (due to interaction effects), then of the inferior competitor (prey species 2) However, virulence effects decrease coexistence when the predator is infected, as they then decrease apparent competition thereby favoring the most competitive species (Fig. 2e). Virulence effects also modulate top-down and bottom-up effects in the system. For instance, when the prey is infected, parasitism incurs a reduction of available energy for higher trophic levels, eventually leading to the loss of the predator (Fig. 2b).

**Figure 2.**
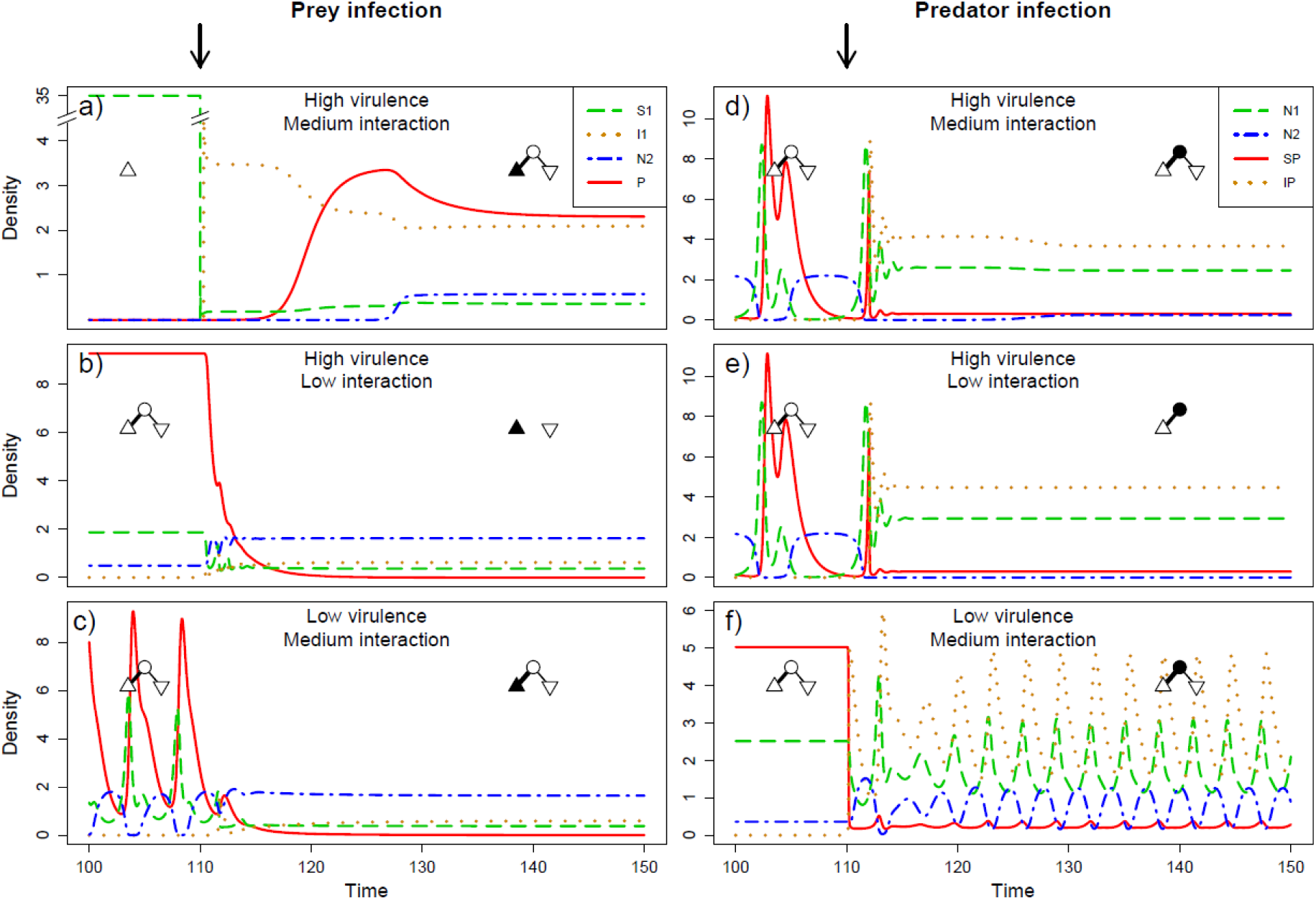
Effects of parasite introduction on coexistence and stability a-c) Prey infection d-f) Predator infection. Arrows indicate the time of parasite introduction. Symbols show the composition of the system before and after adding the parasite: preferred prey 1 (triangle), non-preferred prey 2 (inverted triangle), predator (circle). Infected species are represented in black while non-infected are in white). Prey *N_1_* is shown in green dashed line, prey *N_2_* in blue dashed-dotted line, predator *P* in red solid line, infected prey (a-c) or predator (d-f) individual are shown in orange dotted line. Parameter values: as in Table 1, except a) *f_1_* = 35, *a_1_* = 0.05, *n* = 33, *j* = 1; b) *f_1_* = 15, *a_1_* = 1.2, *n* = 12, *j* = 0.2; c) *f_1_* = 10, *a_1_* = 1.2, *n* = 3.5, *j* = 0.8; d) *a_1_* = 1.2, *m* = 2, *m_i_* = 2.5, *l_i_* = 0.5; e) *a_1_* = 1.2, *m* = 2, *m_i_* = 2.5, *l_i_* = 0.3; f) *a_1_*= 1.2, *m* = 3.2, *m_i_* = 0.5, *l_i_* = 0.5. Note that y-axis of a) is broken.

Interaction effects also act on coexistence. First, they affect the degree of apparent competition among prey species as well as energy availability for higher trophic levels (bottom up effects). For instance, a comparison of Figs 2a and 2b show that for similar virulence effects, predator are not maintained if interaction effects are too weak (Fig. 2b) while larger interaction effects allow such a coexistence (Fig. 2a) by allowing a better energy transfer. Coexistence between the two prey species relies on the balance of direct and apparent competition (Fig. 2a). Too low or too strong interaction effects however lead to the loss of one species, as it changes this balance between the two types of competition (Fig. 2e). These results are coherent with our predictions (Fig. 1b).

Concerning stability, consistent with our predictions (Fig. 1c), we observe that virulence effects do not change stability when the prey is infected (Figs 2a,b), as such effects neither affect the efficiency of energy transfers (interaction rates), nor the distribution of trophic interaction strengths. As expected, virulence effects stabilize the system when predators are infected (Figs 2d,e), as they increase predator mortality. Interaction effects change stability in more complex ways. While in case of prey infection they may stabilize the system by increasing the heterogeneity of interaction strengths (Fig. 2c), in case of predator infection, they may destabilize it by increasing interaction homogeneity or by increasing energy fluxes (Fig. 2f). On Figs 2d-e, we however note that stabilization by virulence effects dominates the complex consequences of interaction effects.

### 3.2. Effects of parasitism in the unstructured model

Now that we have illustrated the possible qualitative effects of parasitism through simulation examples, we vary parasitism continuously in 2D-bifurcation diagrams. We first do so in the unstructured model (Figs 3a,c). X-axis of the bifurcation diagram corresponds to the intensity of virulence effects, while interaction effects are shown on the y-axis. We show variations in composition and stability of the system, depending on the intensity of the effects.

**Figure 3.**
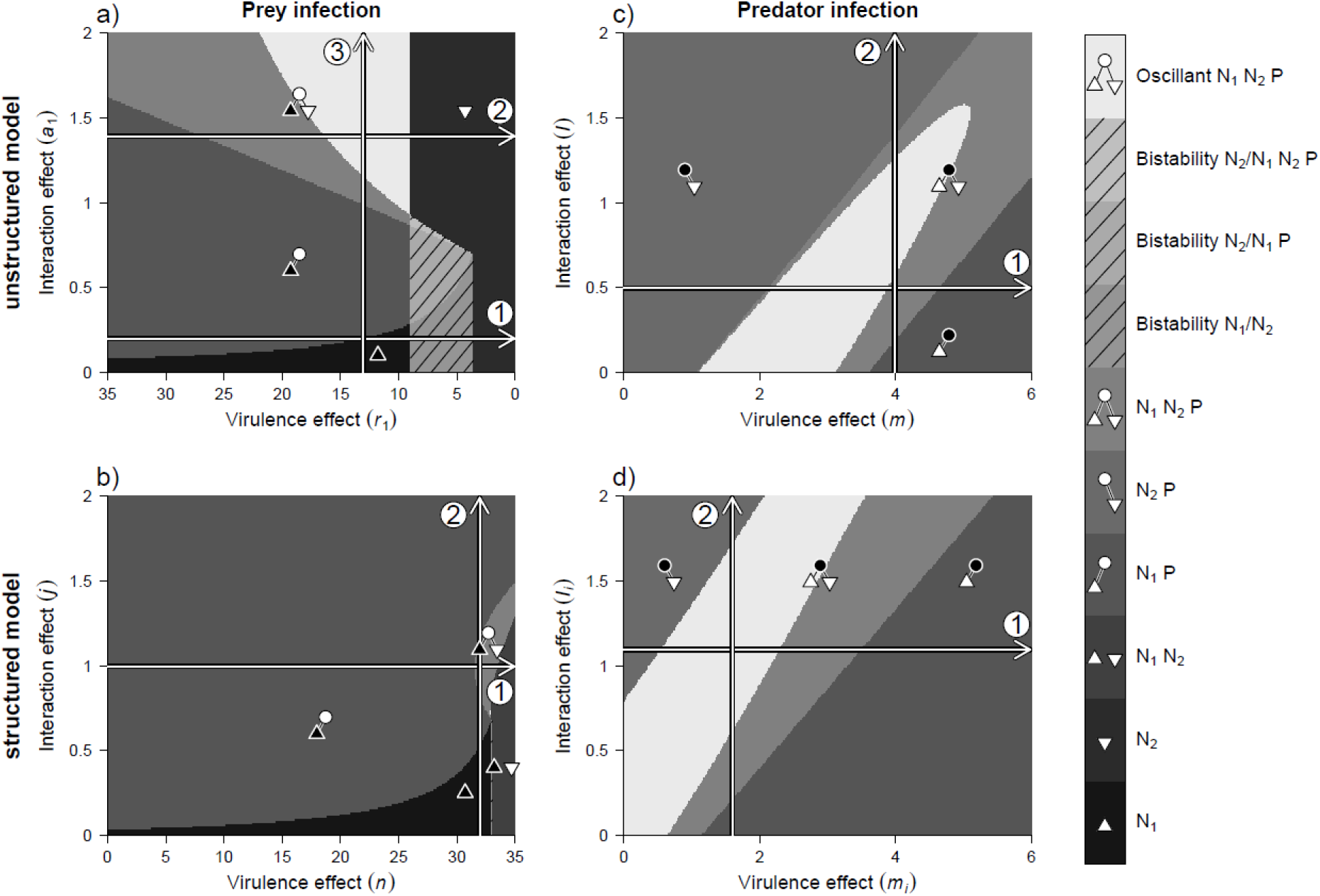
Composition and stability of the system depending on the intensity of virulence (x-axis) and interaction effects (y-axis). a-b) the host is the preferred prey; c-d) the host is the predator. a,c) show results of the unstructured model, b,d) the results of the structured model. Symbols indicate the composition of the system: preferred prey (triangle), non-preferred prey (inverted triangle), predator (circle). Infected species are represented in black. Arrows show the direction of increasing parasite effects: horizontal arrows for the virulence effect and vertical arrows for the interaction effects.

We first analyze how parasitism constrains coexistence. In case of prey infection (Fig. 3a), intermediate virulence effects allow coexistence provided the infected prey undergoes strong predation (arrow 2). Such variations are consistent with our prediction that coexistence requires balance of direct and apparent competition (Fig. 1b). Virulence effects impact coexistence as predicted: at low intensity, we observe a shift in the dominant competitor. Note also that our system exhibits bistability (arrow 1), as expected when interspecific competition dominates intraspecific competition (Case, 2000). Regarding interaction effects, we observe they favor coexistence for the parameter range we consider (arrow 3). Nevertheless, our numerical analyses show that further increases in interaction effects would ultimately lead to a loss of the infected species. Globally, all these results agree with our predictions (Fig. 1b), as parasitism affects coexistence by changing the balance between direct and apparent competition.

Concerning predator infection, our numerical analysis show that prey coexistence is facilitated by intermediate intensities of virulence and interaction effects. This is illustrated on our bifurcation diagram (Fig. 3c, arrow 1 and 2) and in coherence with our predictions (Fig. 1b). A high interaction effect or a small virulence effect (high apparent competition) induces the disappearance of the most consumed and most competitive prey. Contrarily, a small interaction effect or a high virulence (high direct competition) effect leads to loss of the least consumed and least competitive prey.

The intensity of parasitism also affects the dynamical stability of our system (presence/absence of oscillation). For prey infection (Fig. 3a), virulence effects destabilize the system (arrow 2). This contradicts our prediction that stability should not be affected, as virulence effects do not modify interaction strength. A possible hypothesis is that the increase in total prey density (not show) leads to a reduction in competition intensity. Such a decrease in population regulation could be the cause of destabilization. Interaction effects destabilize the system (arrow 3). Such a destabilization may be due to increased energy fluxes from prey to predators (Fig. 1c). In case of predator infection (Fig. 3c), virulence effects and interaction effects destabilize system when they have intermediate intensity. We note that, when increasing simultaneously the two effects, higher stability is eventually achieved.

### 3.3. Coexistence and stability in structured models of infection

We similarly analyzed the structured models of prey and predator infection (Figs 3b,d). Coexistence is favored at intermediate intensity of virulence and interaction effects regardless of the infected species. Thus, results on coexistence remain consistent with our predictions (Fig. 1b) and with the results observed for the unstructured model. Some finer differences however exist between the structured and unstructured models of prey infection (Fig. 3a vs Fig. 3b). Virulence effects still lead to a reduced competitive ability of the infected species, which favors the co-occurrence of the two competitors (arrow 1). However, in the structured model, the explicit dynamics of parasite (susceptible-infected) provides an additional feedback loop between predator and parasite effects acting on prey species 1. Therefore, global negative pressures on this species are more balanced and its competitive ability is less reduced compared to the unstructured scenario. This species remains dominant for a larger set of parameters, which reduced the possibility of coexistence. Hence, the coexistence area appears highly reduced compared to the unstructured model. The explicit dynamics of the parasite in the structured model also lead to a competition between predator and parasite populations (Fig 4). Virulence effects favor the parasite in this competition. They lead to an increase in density of infected and to a decrease in predator density (Fig 4, arrow 1). Contrarily, interaction effects ultimately favor the predators. Larger effects then decrease parasite density while increasing predator density (Fig 4, arrows 2 and 3). Consequently, while intermediate effects of parasitism are still required to maintain coexistence, the mechanism now relies not only on the balance between direct vs apparent competition among prey species, but also on a balanced competition between predators and parasites. This competition may lead to the disappearance of parasites at large interaction effects (Figs 4, S1), and of the predator at large virulence effects (Figs 3a-b, 4, S1). In scenarios of predator infection, structured and unstructured models give qualitatively similar results (Fig. 3c vs Fig. 3d).

**Figure 4.**
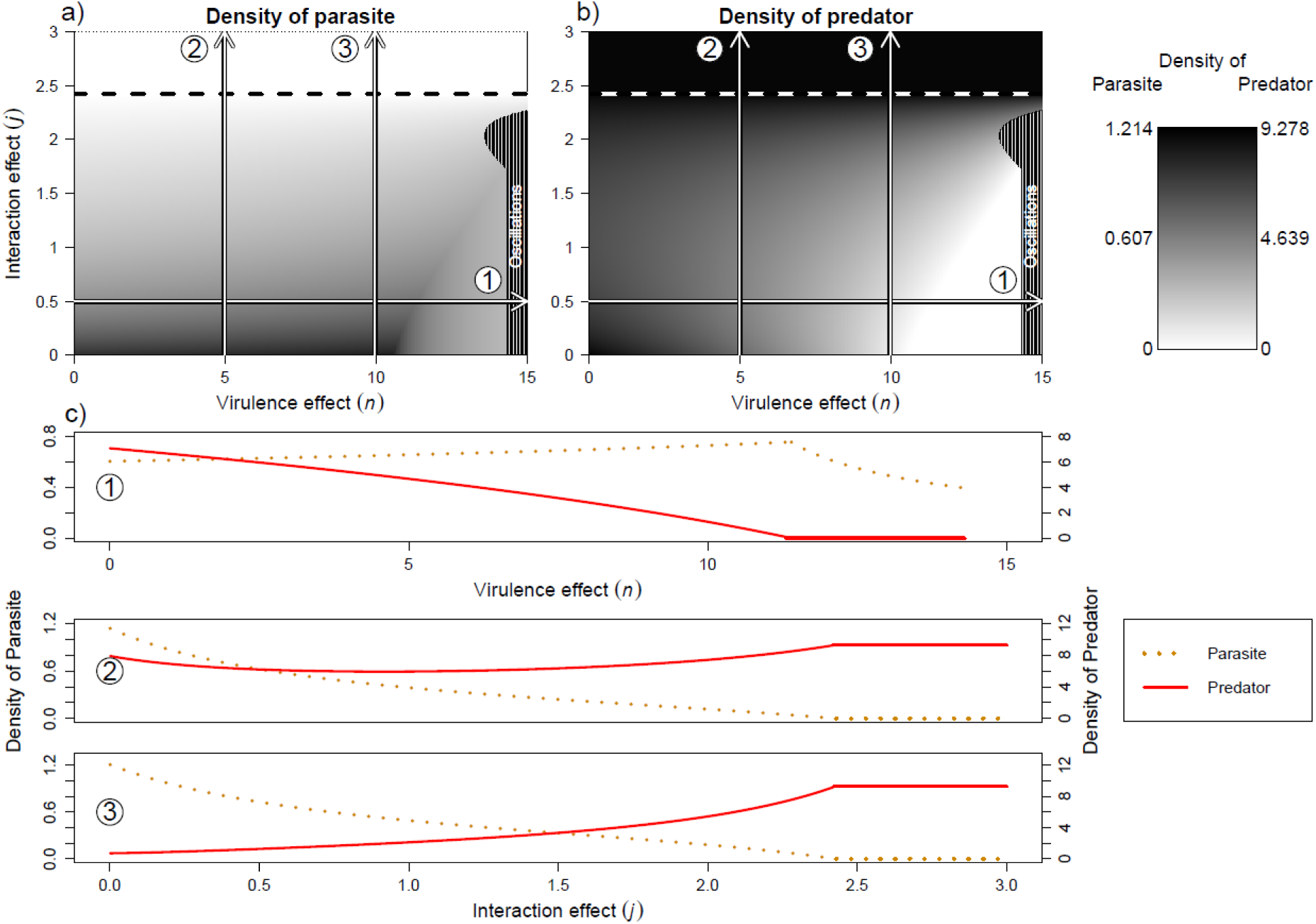
Analysis of predator-parasite competition. Levels of prey infection (a) (proxy for parasite density) and of predator density (b) depending on the intensity of virulence (x-axis) and interaction effects (y-axis). Arrows show the direction of increasing parasite effects (arrow 1 shows increased virulence effects, arrows 2 & 3, increased interaction effects). Above the dashed lines, parasite cannot persist. The right black area corresponds to oscillating systems. c) Three bifurcation diagrams, corresponding of the three arrows, showing the contrasting effects of virulence effects (that favor the parasite population) and of interaction effects (that favor the predator population). (1) *j* = 0.5, (2) *n* = 5 and (3) *n* = 10. The lines show predator density (red solid lines) and infection levels (orange dotted line).

Effects of parasitism on stability are more idiosyncratic. For infected prey (Fig. 3b), the area of oscillations is greatly reduced in the structured system. This increased stability of structured prey infection system (compared to the non-structured case) is commonly observed for various sets of parameters (Fig. A1). When the parasite is maintained in the system, virulence and interaction effects seldom lead to an oscillating system (Fig. A1a-b) or stabilize an unstable one (Fig. A1c). Such stabilizing effects may be explained by the fact that the structured model explicitly accounts for an additional negative feedback between the predator and parasite populations. Regarding predator infection (Fig. 3c), virulence effects stabilize the coexistent system (arrow 1), as predicted (Fig. 1c). Interaction effects (arrow 2) first destabilize the system, in coherence with our predictions (Fig. 1c). Further increases in interaction effects eventually lead to the destabilization of the module and to the loss of the prey species 1 (preferred by the predator).

## 4. Discussion

The present work uses simple models to highlight and understand mechanistically possible consequences of parasitism for coexistence and stability in predator-prey systems. We show that such consequences depend on the type of parasitism (virulence vs interaction effects), on the species that is infected (predator or prey), but that they can be understood to some extent based on classical ecological theories based on apparent competition (Holt, 1977) and interaction strength (McCann et al., 1998; Rip & McCann, 2011). More precisely, parasites affect coexistence within trophic level by changing the balance between direct and apparent competition. In the case of infected prey, parasites also modify coexistence among trophic levels by altering bottom-up effects and through competition with predators. Assessing the latter effect however requires the construction of structured model that allow explicit variations in the parasite populations. While parasitism can affect stability positively or negatively depending on the scenario, the ratio between energy fluxes and predator mortality rates largely explain the effects of predator infection on system stability, as proposed in previous works (Rip & McCann, 2011; Rosenzweig, 1971).

Within a given trophic level (among prey species), we find that parasites may alter coexistence by affecting the relative intensity of direct and apparent competition. First, parasites may reduce its host competitive ability (for instance through virulence effects), allowing coexistence with an inferior competitor (“parasite-mediated competition” *sensu* Hatcher & Dunn [2011]). Such a mechanism of coexistence is coherent with previous theoretical works (e.g.Anderson & May [1986]) and has also been observed in experiments and field investigations (Callaway & Pennings, 1998; Decaestecker et al., 2015; Kiesecker & Blaustein, 1999; Park, 1948; Price, Westoby, & Rice, 1988; Schall, 1992). For instance, Schall (1992) studied the coexistence of lizard species in Caribbean islands and observed that *Anolis wattsi* was present only when *A. gingivinus* was infected by the malarial parasite *Plasmodium azurophylum,* a parasite that has clear virulence effects.

Parasites infecting predators also affect coexistence and biomass distribution among trophic levels as they reduce top-down effects and alter apparent competition between prey species. By increasing predator mortality (e.g. through virulence effects), such parasites act as a top-predator (Wilmers, Post, Peterson, & Vucetich, 2006) and may induce trophic cascades that increase prey density. Such parasite-mediated trophic cascades have been observed in nature (Buck & Ripple, 2017). For instance, Lindström et al. (1994) show that the infection of red foxes by *Sarcoptes scabiel* can lead to increased hare and grouse densities. However, such positive consequences of predator infection on prey abundances can be redistributed asymmetrically among prey species, as such parasites also alter apparent competition. Given virulence effects, the parasites of predators would lead to a release of apparent competition as they decrease predator populations. Contrarily, parasites incurring interaction effects may reinforce apparent competition. Such modifications of prey composition have been observed for both effects. Empirical studies showed that virulence effects reduce top-down control and affect prey composition (Dobson & Crawley, 1994; Hartley, Detling, & Savage, 2009). Other experiments also showed that increased predation rate due to parasitism of the predator (i.e., interaction effects) can lead to a shift in the dominant prey species. Bernot & Lamberti (2008) for instance found that snails (*Physa acuta*) infected by a trematode (*Posthodiplostomum minimum*) have a greater grazing rate leading to a periphyton community dominated by *Cladophora* whereas, without parasite, periphyton is dominated by diatom and blue-green algae. Furthermore, system where infection reduce grazing rate presents similar results (Wood et al., 2007).

Parasites of prey species also affect higher trophic levels through bottom-up effects. They have a particular interaction with the predators of their host for which they become both a prey and a competitor (Sieber & Hilker, 2011), which is close to intraguild predation. Most previous studies focused on the effects of predators on infection levels. With this point of view (i.e. focusing on the effects of predators on parasites), Packer, Holt, Hudson, Lafferty, & Dobson (2003) developed the “healthy herd hypothesis” observed in theoretical and empirical works (Anderson & May, 1986; Duffy et al., 2005; Lafferty, 2004). By reducing host/prey density below a threshold, predators lead to parasite extinction. This effect increases when predator consumes preferentially infected prey (Hethcote, Wang, Han, & Ma, 2004; Packer et al., 2003), for instance due to interaction effects. Such observations are consistent with our results. In our model, interaction effects systematically favor predators in their competition with parasites. Such outcomes have also been observed experimentally. As shown by Duffy et al. (2005), the prevalence of parasite *Spirobacillus cienkowskii* in *Daphnia dentifera* population decreases when the abundance of bluegills (a predator of daphnia) increases. Our work also clarifies conditions under which parasites have competitive or facilitative effects on predators, through modifications of bottom-up effects. When parasites have interaction effects, they allow the persistence of predators by making prey more available (facilitative effect). Such effects are consistent with previous theoretical works (Hethcote et al., 2004). However, when parasites have mostly virulence effects, their negative impact on prey density may lead to the disappearance of the predator (competitive effect). Such competitive effects are consistent with earlier theoretical works (e.g. Anderson & May [1986]). Parasites and predators then interact as competitors, sharing a common resource: the prey species. Such a competition ultimately reduces coexistence. Empirical examples of such dynamics exist. Banerji et al. (2015) showed experimentally that, by reducing prey density (*Paramecium caudatum*) parasites may lead to a reduction of predator density (*Didinium nasutum*).

Effects of parasites on stability seem to be highly context-dependent (Lafferty et al., 2008; Wood & Johnson, 2015). Previous theoretical and empirical studies report stabilizing effects through regulation of host populations (Anderson & May, 1978; Cáceres et al., 2014; Hilker & Schmitz, 2008) or through parasite-mediated coexistence (Dobson, 2004). Ong & Vandermeer (2015) for instance showed that adding not only predators, but also parasites allows for a more stable biological control. Other studies report destabilizing effects (Anderson & May, 1978, 1986; Grenfell, 1992; Hudson, Dobson, & Newborn, 1998; May & Anderson, 1978), for instance due to increased vulnerability to predation (Ives & Murray, 1997), or when parasites create time lags in dynamics (Hudson et al., 1998; May & Anderson, 1978). In our model, stability outcomes are equally variable, as parasites may stabilize an unstable system or destabilize a stable one, even within a given parasitism scenario, depending on the considered set of parameters. Nevertheless, our model highlights how some of these results on stabilization/destabilization can be related to general theories of stability in consumer-resource interactions. We systematically assessed two basic hypotheses: that heterogeneity in interaction strengths increases stability (McCann et al., 1998) and that stability depends on relative energy fluxes (ratio between attack rates and predator mortality rates [Rip & McCann, 2011; Rosenzweig, 1971]). Our model shows that the second hypothesis largely explains the patterns we observe in case of predator infection. We indeed observe that parasites of predators have a stabilizing effect in case of virulence effects (that increase in predator mortality), but a destabilizing effect when they induce interaction effects (that increase of attack rate). The stabilizing effects of virulent parasites infecting predators are consistent with previous theoretical results (Hilker & Schmitz, 2008) while the destabilization due to interaction effects had also been observed in model by Bairagi & Adak (2015).

While we categorized the effects of parasites in two schemes – virulence and interaction effects, most parasites are likely to alter simultaneously life-history traits (with consequences for mortality and/or reproduction) and species behavior or physiology (with consequences for interaction strength). Interestingly, in some scenarios, coexistence can only be reached when combining the two effects. For instance, in prey infection scenarios, combined effects favor coexistence. When considering predator infection, simultaneous increases in both effects allow the maintenance of the whole system, while an increase in only one of the two effects ultimately reduces species coexistence. While early studies of parasitism focused on virulence effects (Holt & Pickering, 1985; Park, 1948), modifications of trophic interactions (i.e. interaction effects) have been investigated in trophically-transmitted parasites within the framework of the manipulation hypothesis (Bethel & Holmes, 1977; Poulin & Maure, 2015). In this context, manipulative parasites can increase host vulnerability and thereby facilitate their own transmission probability to next host (Cézilly et al., 2010). However, interaction effects do not require host manipulation and may actually emerge more generally as by-products of physiological changes incurred by the infection of either prey (Duffy et al., 2005; Hudson et al., 1992; Peterson & Page, 1988) or predator species (Arnott, Barber, & Huntingford, 2000; Dick et al., 2010; Wright, Wootton, & Barber, 2006). Virulent parasites lead by definition to modifications of host energy requirement or allocation (Hall, Becker, & Cáceres, 2007). To face such energetic challenges, hosts may reduce or increase their activity, thereby affecting the probability of encounters with predators or prey. As a result, infection may increase vulnerability to predation (Gehman & Byers, 2017; Peterson & Page, 1988) or reduce the predation rate of consumer species (Coop, Sykes, & Angus, 1982; Wood et al., 2007). Alternatively, to compensate energy reduction due to parasites, some host species increase their predation activities (Khokhlova, Krasnov, Kam, Burdelova, & Degen, 2002; Lettini & Sukhdeo, 2010). All these studies are mostly focused on a given trophic interaction, and the effects of virulence and interaction effects at the community level remain understudied. Banerji et al. (2015) have however analyzed a tri-trophic food chain with a resource, a consumer, a predator and a parasite of the consumer. They showed that infection leads to variations in growth rate (implying virulence effects), changes in consumption rate (thus interaction effects), with implications for the dynamics of each species. In their case, infection of a *Paramecium* decreased its growth rate and its cell size, increased its velocity and its grazing rate, but did not modify its vulnerability.

While we mostly focus here on the effects of parasitism on the ecological dynamics of the community, we propose that parasitism may affect the evolution of predator foraging activities. According to the theories of optimal and adaptive foraging, selection favors predators foraging on the energetically most profitable prey (Charnov, 1976a, 1976b; Emlen, 1966; MacArthur & Pianka, 1966). Prey profitability being defined as the ratio between energy content of the prey and its handling time for a given search time, we note that all these components are likely to be modified by parasites. Parasites altering the vulnerability of host species reduce either the search time or the handling time. Virulent parasites, when reducing host density should increase search time. Parasites are also able to increase (Hall et al., 2007) or decrease (Caddigan, Pfenning, & Sparkes, 2017; Flick, Acevedo, & Elderd, 2016; Forshay, Johnson, Stock, Peñalva, & Dodson, 2008) the energy content of their host. We therefore expect that the adaptation of predator foraging in response to parasitism may lead to vast changes in the relative intensity of trophic interactions, thereby altering coexistence conditions (direct modifications of apparent competition, Holt [1977]) or system stability (by altering the distribution of interaction strengths, McCann et al. [1998]). Such eco-evolutionary dynamics thus offer important perspectives to better understand the effects of parasitism on ecological networks.

## Acknowledgements

The authors thank D. Claessen and T. Spataro for helpful discussions.

## Authors’ contributions

LP, VM and NL designed the model and the computational framework and analyzed the data. LP performed analytic calculations and performed the numerical simulations. LP wrote the first draft, NL edited the first draft and all authors participated to further rewriting of the manuscript.

## Supplementary

### S1 Composition and stability of prey-infected system for various set of parameters

**Figure S1.**
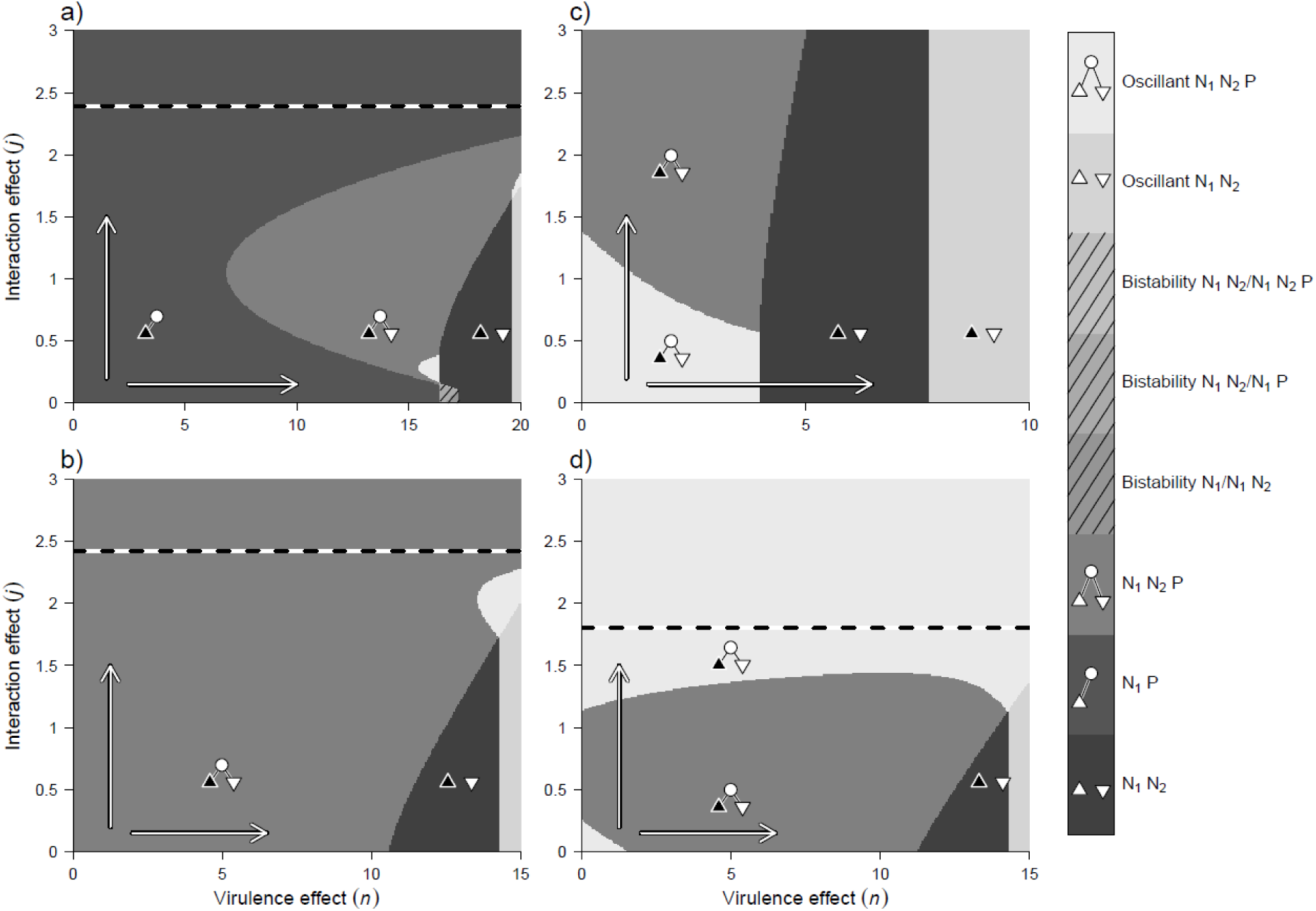
Variations in the composition and stability of the system given prey infection, for various initial compositions of the uninfected system. Symbols indicate the composition of the system: preferred prey (triangle), non-preferred prey (inverted triangle), predator (circle). Host species are represented in black. Arrows show the direction of increasing parasite effects. Horizontal dashed lines correspond to limits above which the parasite cannot persist. Parameter values: as in Table 1 except a) *f*_*1*_ = 20, *a*_*1*_ = 0.6; b) *f*_*1*_ = 15, *a*_*1*_ = 1.2; c) *f*_*1*_ = 10, *a*_*1*_ = 1.2; d) *f*_*1*_ = 15, *a*_*1*_ = 1.5.

